# Airway bacterial and fungal microbiome in chronic obstructive pulmonary disease

**DOI:** 10.1101/2020.10.05.327536

**Authors:** Haiyue Liu, Zhenyu Liang, Nannan Cao, Xilan Tan, Zuheng Liu, Fengyan Wang, Yuqiong Yang, Chunxi Li, Yan He, Jin Su, Rongchang Chen, Zhang Wang, Hongwei Zhou

## Abstract

**Background:** Little is known about airway mycobiome, and its relationship with bacterial microbiome in chronic obstructive pulmonary disease (COPD).

**Methods:** Here we report the first simultaneous characterization of sputum bacterial and fungal microbiome in 84 stable COPD and 29 healthy subjects, using 16S ribosomal DNA and fungal internal transcribed spacer DNA sequencing.

**Results:** Ascomycota predominated over Basidiomycota in fungal microbiome both in COPD patients and healthy controls. *Meyerozyma*, *Candida*, *Aspergillus* and *Schizophyllum* were most abundant at the genus level. There was a significant inverse correlation between bacterial and fungal microbial diversity, both of which altered in opposite directions in COPD patients versus controls, and in frequent versus non-frequent exacerbators. An enhanced bacterial-fungal ecological interaction was observed in COPD patients, which was characterized by higher proportion of co-occurrence intrakingdom interactions and co-exclusive interkingdom interactions. In COPD, four mutually co-occurring fungal operational taxonomic units (OTUs) in *Candida palmioleophila*, *Aspergillus* and Sordariomycetes exhibited co-exclusive relationships with other fungal OTUs, which was specifically present in frequent exacerbators but not in non-frequent exacerbators. Conversely, the mutual co-occurrence interactions between bacterial OTUs in *Rothia mucilaginosa*, *Streptococcus*, *Veillonella* and *Prevotella*, showed up in non-frequent exacerbators but not in frequent exacerbators. The perturbed bacterial-fungal interactions in COPD were associated with increased airway inflammatory mediators such as IL-6 and IL-8.

**Interpretation:** The disruption of airway bacterial-fungal community balance, characterized by the loss of commensal bacterial taxa and enriched pathogenic fungal taxa, is implicated in COPD. The airway mycobiome is an important cofactor mediating COPD pathogenic infection and host inflammation.

**Clinical Trial Registration:** www.clinicaltrials.gov (NCT 03240315).

## Introduction

Chronic obstructive pulmonary disease (COPD) is characterized by chronic airway inflammation resulting in irreversible decline in respiratory function and capacity. Bacterial, fungal and viral infections drive airway inflammation, and are associated with poorer disease outcome(1) and declined lung function(2, 3). The airway microbiome, the collective airway microbial community, is hypothesized to mediate the interactions between pathogenic infection and host inflammatory response(4, 5). Through interacting with bacteria and mucosal immune system(6), the fungal community can be a cofactor for airway inflammation and COPD progression(7). Essentially all previous airway microbiome studies, however, have focused on bacterial community in COPD(4, 8–14). The non-bacterial members of airway microbiome in particular the fungal microbiome (or mycobiome), despite being of clinical relevance, have been largely underappreciated(7). Recent studies have reported the airway fungal composition in asthma(15), bronchiectasis(16), cystic fibrosis(17, 18), HIV(19) and lung transplantation(20); however, little is known on the mycobiome in COPD. The ecological interaction between airway bacterial and fungal microbiome and its role in COPD pathogenesis remains unexplored.

Susceptibility to frequent exacerbations represents an independent clinical phenotype in COPD, the ‘frequent exacerbator’ phenotype, and is associated with poorer clinical outcome(21, 22). The Global Initiative for Chronic Obstructive Lung Disease (GOLD) 2019 has redefined the measure of disease severity to recognized the high exacerbation risk(23) (>=2 exacerbations and /or 1 hospitalization in the previous year). The pathophysiology underlying the frequent exacerbation phenotype is manifested by an interplay between enhanced airway immune responses, bacterial and fungal colonization and dynamic lung hyperinflation, that together predispose patients to persistent inflammation and recurrent exacerbations(22). Identifying markers that predict patient exacerbation frequency is of great importance for COPD management. Difference in baseline respiratory microbiome composition was hypothesized to explain the different exacerbation frequency in COPD patients(21). However, studies assessing baseline airway microbiome have not found significant differences in bacterial composition between frequent and non-frequent exacerbators(4, 14). Recent longitudinal studies suggested that temporal variability of the sputum microbiome could be associated with COPD exacerbation frequency(14, 24). However, measuring microbial temporal variability requires serial sampling of sputum microbiome in multiple timepoints and is therefore not clinically practical. Assessing the airway fungal microbiome might open up opportunities in identifying novel markers for the frequent exacerbator phenotype.

Here we characterized the airway bacterial and fungal microbiome simultaneously in clinically stable COPD patients. We hypothesize that the ecological interactions between bacterial, fungal microbiome and host inflammation are associated with disease and exacerbation frequency. We showed that bacterial and fungal microbiome co-altered in COPD. The perturbation of bacterial-fungal interactions in COPD was associated with host inflammation and the frequent exacerbator phenotype.

## Materials and Methods

### Subjects and samples

Sputum samples of 113 individuals, including 84 stable COPD patients and 29 healthy controls, were collected in the First Affiliated Hospital of Guangzhou Medical University. COPD patients were divided into frequent (FE: exacerbation events >=2 or 1 hospitalization due to exacerbation of COPD/past year) and non-frequent exacerbators (NE). The study was approved by the ethics committee of the First Affiliated Hospital of Guangzhou Medical University and was registered in www.clinicaltrials.gov (NCT 03240315). All subjects provided written informed consent in accordance with the Declaration of Helsinki.

### Bacterial and fungal microbiome sequencing

Bacterial genomic DNA was extracted from selected sputum plugs using a Total Nucleic Acid Extraction Kit (Bioeasy Technology, Inc., Shenzhen, China) as per the manufacturer’s instructions. Negative controls for extraction (no sputum) and PCR amplification (no DNA template, ddH_2_O only) were included in each experiment. The extraction negative controls were subsequently sequenced to identify any potential contaminating bacterial/fungal species. The V4 hypervariable region of bacterial 16S rRNA gene and fungal 18S–28S rRNA gene internally transcribed spacer region ITS1 DNA were amplified using barcoded primers, and were sequenced using iTorrent sequencing platform.

### Sequence processing and analysis

Sequence processing and analysis were performed using QIIME 1.9.1(25). The obtained sequences were de-multiplexed, trimmed of barcodes and primers, and filtered if they contained ambiguous bases or mismatches in the primer regions, according to the BIPES protocol(26). Chimeras were filtered out using UCHIME in *de novo* mode(27). After quality filtering and chimera removal, 16S rRNA gene sequencing resulted in a median read depth of 4,380, ITS1 DNA sequencing resulted in a median read depth of 9,783. Both 16S rRNA V4 region and ITS1 DNA sequencing data of all subjects were subsampled to a uniform depth of 1,000 reads based on rarefaction curve asymptotes and Good’s coverage values. Comparable rarefaction depth has been used in airway microbiome analyses(4, 28).

High-quality sequence reads for bacterial 16S rRNA V4 and fungal ITS1 region were clustered into operational taxonomic units (OTUs) using USEARCH v11(29) in de novo mode with 97% sequence similarity cutoff. The taxonomy of representative 16S rRNA gene sequences were determined using PyNAST with the Greengenes 13_8 database as reference(30). The taxonomy of representative ITS sequences were determined using the QIIME_ITS database as reference (version information: sh_qiime_release_s_28.06.2017)(25). The taxonomy of highly abundant unclassified fungal OTUs was further determined using the phylotyping algorithm in MEGAN5 (http://ab.inf.uni-tuebingen.de/software/megan5/)(31). Briefly, the OTU representative sequence was BLASTn-searched (BLAST v2.5.0) against the non-redundant reference database and the last common taxonomic rank of all sequence hits with >97% was assigned to that OTU. OTUs with >0.5% average abundance were selected for downstream analysis. The sequences were deposited in the European Nucleotide Archive (ENA) under accession numbers PRJEB27507.

The biome data were filtered using the filter_otus_from_otu_table.py script with the parameter (-s 3) to remove low-abundance OTUs. Fifty-four OTUs (28 bacterial and 16 fungal OTUs) were selected that were >0.5% average abundance for the major of OTU-based analysis. Airway microbial alpha diversity (diversity within samples) was calculated using the Shannon indices. Airway microbial beta diversity (composition dissimilarity between samples) was determined by using the unweighted UniFrac distance and visualized in Principal Coordinate Analysis (PCoA). Adonis was used to estimate statistical significance. Differential features between groups were identified using a linear discriminant analysis (LDA) effect size (LEfSe) method with a threshold of logarithmic LDA score 2.0(32). Random forest analysis was performed using the OTU data selected by LEfSe using Weka 3.8 (https://www.cs.waikato.ac.nz/ml/weka/) with a 7-fold cross-validation(33). Co-occurrence and co-exclusion relationships between the 54 abundant bacterial and fungal OTUs were estimated using SparCC algorithm(34), known for its robustness to compositional effects in microbiome dataset. The p-value was estimated by 100 bootstraps and the correlations with p<0.05 were retained. Association between bacterial and fungal OTUs and inflammatory mediators was assessed using Spearman correlation. Network was visualized using Cytoscape 3.6.0 (https://cytoscape.org/)(35). The false discovery rate (FDR)(36) method was used to adjust *P*-values for multiple testing wherever applicable.

## Results

### Overview of sputum bacterial and fungal microbiome

Sputum samples were collected from 84 stable COPD patients and 27 healthy controls (Table 1, Table S1). Consistent with previous studies(4, 8–11, 14, 24, 37), the majority of bacterial taxa belongs to Firmicutes (28.3%), Bacteroidetes (27.2%), Proteobacteria (24.6%), Actinobacteria (7.0%) and Fusobacteria (5.7%) at the phylum level. For the fungal microbiome, 76.6% sequences belong to Ascomycota (71.7%) or Basidiomycota (4.9%) and 17.7% sequences to unidentified fungi taxa. The most abundant fungal genera are *Meyerozyma*, *Candida*, *Aspergillus* and *Schizophyllum* (>1% average) (Fig. 1). There was a significant negative correlation between bacterial and fungal alpha diversity (Shannon, Spearman’s rho=−0.172, p=0.04). No significant association was found between bacterial or fungal microbiome with age, gender, smoking status and predicted FEV1% (Fig. 1).

**TABLE 1.** Major clinical characteristics of subjects.

|  | Healthy<br>(n=29) | COPD<br>(n=84) | Historical exacerbation<br>frequency |  |
| --- | --- | --- | --- | --- |
|  |  |  | NE group<br>(n=52) | FE group<br>(n=27) |
| Age, mean (SD) | 44.28<br>(23.10) | 64.55<br>(8.83)*** | 66.02 (7.72) | 61.50 (9.76) |
| Gender,<br>n(male/female) | 21/8 | 81/3*** | 50/2 | 25/1 |
| Smoking, n(yes/no) | 9/20 | 74/7*** | 45/7 | 23/0 |
| FEV1% predicted,<br>mean (SD) | NA | 51.11 (23.46) | 55.63 (23.54) | 41.71 (19.91)* |
| ICS+LABA, n(yes/no) | NA | 51/33 | 34/18 | 17/9 |

FEV1: forced expiratory volume in 1s. ICS+LABA: Combination of inhaled corticosteroid and long-acting bronchodilators. \*\*\*p<0.001, \*\*p<0.01, \*p<0.05

**Figure 1.**
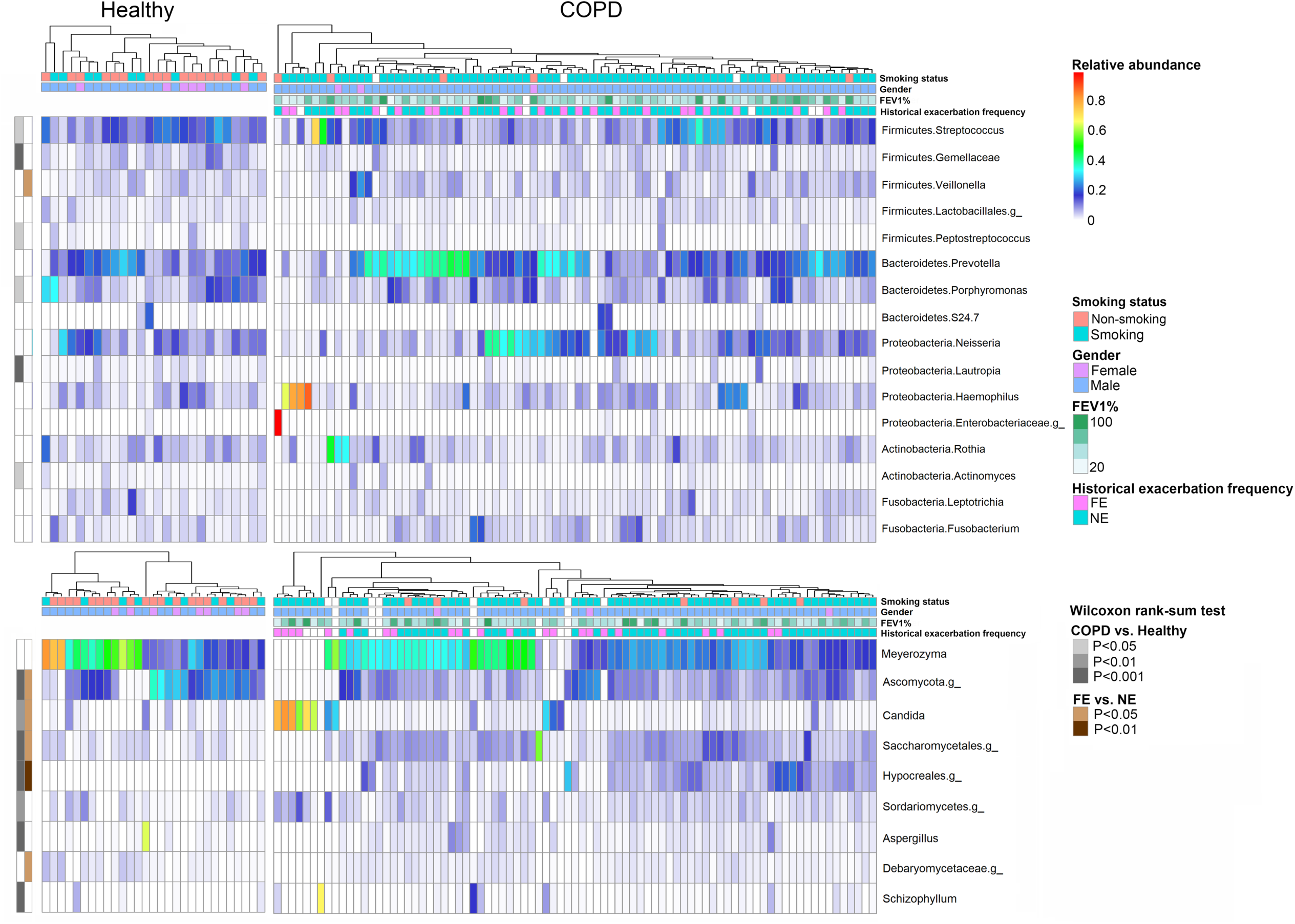
Heatmaps showing the major bacteria and fungi genera (>1% average abundance) in COPD patients and healthy controls. Each column represents an individual grouped first by healthy or COPD subjects and then clustered by bacterial or fungal microbiota composition. The rows on the top represent demographic factors.

### Sputum bacterial and fungal microbiome in COPD patients and healthy controls

The bacterial composition shifted in COPD patients compared to healthy subjects, with a slight decrease in alpha diversity (Fig. 2a) and a significant decreased abundance of genera *Streptococcus*, *Peptostrptococcus*, *Porphyromonas*, *Lautropia* and *Actinomyces* (Wilcoxon, FDR p<0.05, Fig. 1). Conversely, the fungal diversity significantly increased in COPD (Fig. 2b), with a significant increased abundance in *Candida* and *Schizophyllum*, and three unclassified genera in Sordariomycetes, Saccharomycetales and Hypocreales (Wilcoxon, FDR p<0.05, Fig. 1). Beta-diversity analysis indicated a better separation between healthy and COPD groups using fungal than bacterial composition (Adonis, bacteria: R^2^=0.016, p<0.01; fungi: R^2^=0.061, p<0.01, Fig. 2a-b). LEfSe analysis identified seven bacterial OTUs (bOTUs) and seven fungal OTUs (fOTUs) associated with disease state (LDA>2.0, Fig. S1-2). Random forest analysis discriminated COPD patients from controls with an area under the curve (AUC) of 0.83, 0.91 and 0.97 using these bacterial, fungal and their combined OTUs, respectively (Fig. 2c). Sub-analysis using 52 age, gender and smoking-status matched subjects indicated that the observed microbiome differences were not related to these factors (Table S2, Fig. S3).

**Figure 2.**
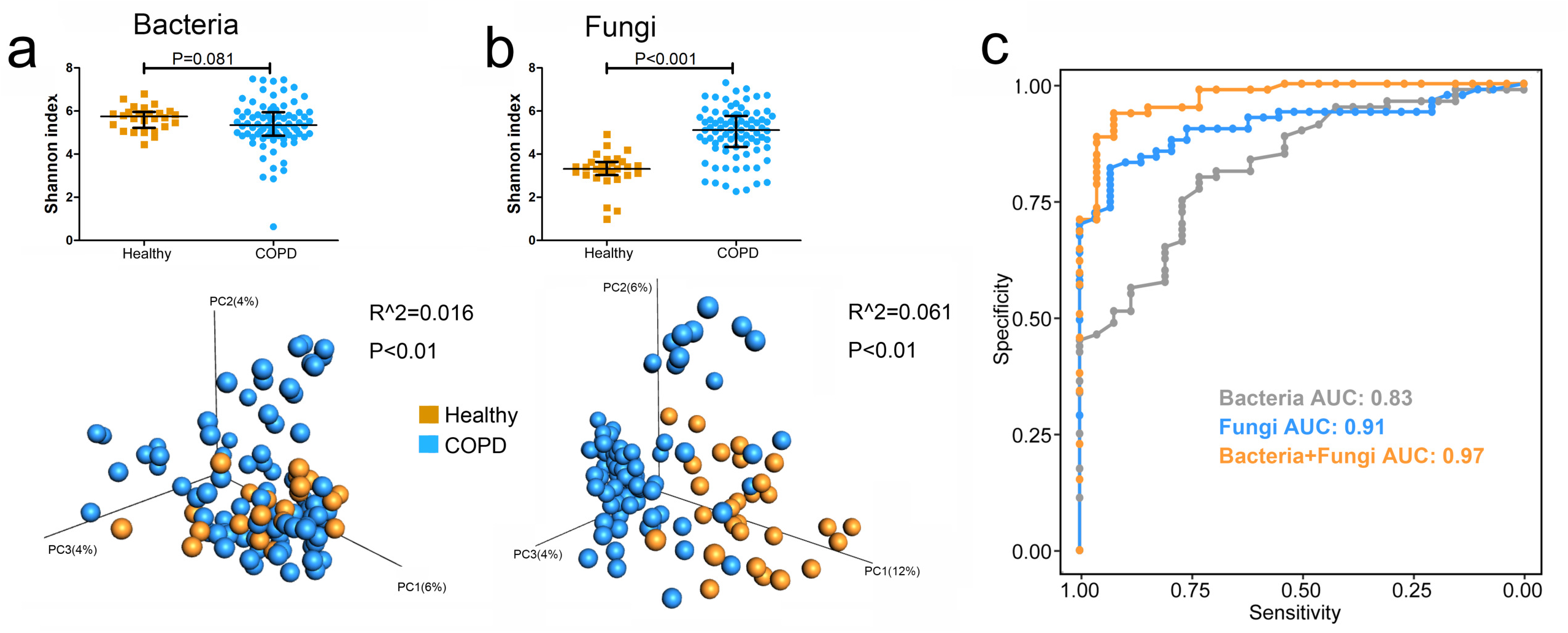
Airway bacterial and fungal composition between COPD patients and healthy controls. Shannon index for **(a)** bacterial and **(b)** fungal microbiome in COPD patients and healthy controls. Beta diversity was assessed based on unweighted UniFrac distances and plotted in PCoA. **(c)** The AUC curves for the random forest models in separating COPD and healthy groups using the LEfSe-selected bacterial, fungal and their combined OTUs.

### Sputum bacterial and fungal microbiome in frequent and non-frequent exacerbators

COPD patients were divided into frequent (FE: exacerbation events ≥2 or 1 hospitalization due to exacerbation of COPD/past year) and non-frequent exacerbators (NE). Patient demographic factors are overall comparable between the two groups except for a significantly higher FEV1% predicted in NE group (Table 1). Bacterial alpha diversity was significantly higher (Fig. 3a) in FE compared to NE group, whereas fungal alpha diversity showed the opposite trend (Fig. 3b). *Veillonella* was significantly decreased in FE group (Wilcoxon, FDR p<0.05, Fig. 1), whereas fungal genera *Candida* was significantly increased. Beta-diversity analysis also indicated a better separation between FE and NE for fungal compared to bacterial composition (Adonis, bacteria: R^2^=0.019, p<0.01; fungi: R^2^=0.046, p<0.01, Fig. 3a-b). Seven bOTUs and seven fOTUs were associated with exacerbation frequency using LEfSe (LDA>2.0, Fig. S4-5). Random forest analysis showed an AUC value of 0.78, 0.74 and 0.81 in separating the two groups using these bacterial, fungal and their combined OTUs respectively (Fig. 3c).

**Figure 3.**
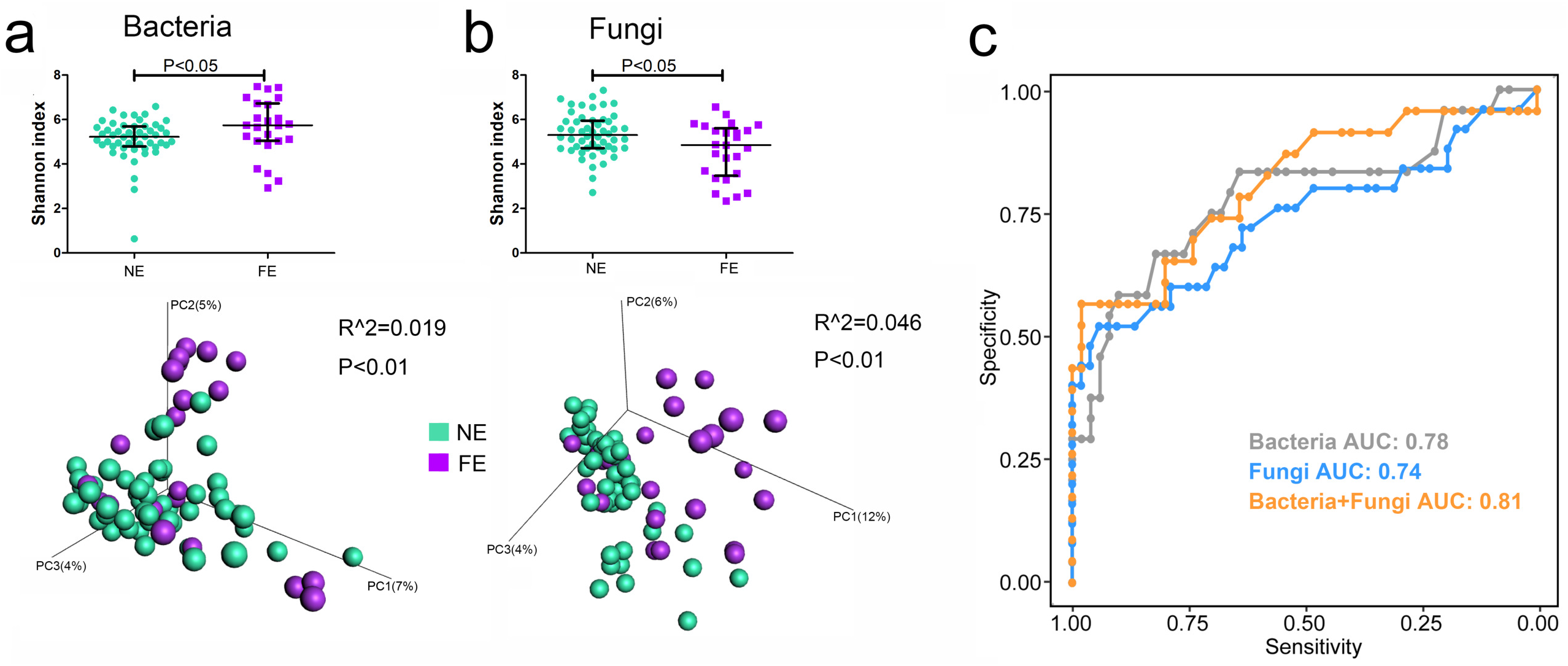
Airway bacterial and fungal composition between COPD frequent and non-frequent exacerbators. Shannon index for **(a)** bacterial and **(b)** fungal microbiome in COPD frequent (FE) and non-frequent exacerbators (NE). Beta diversity was assessed based on unweighted UniFrac distances and plotted in PCoA. **(c)** The AUC curves for the random forest models in separating FE and NE using the LEfSe-selected bacterial, fungal and their combined OTUs.

### Bacterial-fungal interactions in COPD patients and healthy controls

To explore ecological interactions between airway bacterial and fungal microbiome, we performed network analyses using 54 bacterial and fungal OTUs (>0.5% average relative abundance) using the SparCC algorithm(34). We observed considerable differences in bacterial-fungal interactions between COPD patients and controls. For COPD patients, 244 significant correlation pairs comprising of 144 bacteria-bacteria (B-B), 32 fungi-fungi (F-F) and 68 bacteria-fungi (B-F) interactions were identified (Fig. 4a-b, Fig. S6, p<0.05). Among them, 100 (69.4%) B-B, 23 (71.9%) F-F and 21 (30.9%) B-F correlations were positive, indicating predominant co-occurring intrakingdom interactions and co-exclusive interkingdom interactions in COPD (Fig. 4a). Among the inverse relationships between bacterial and fungal OTUs, the correlations between bOTU2 *Prevotella melaninogenica* and fOTU15 *Leucosporidium scottii*, and between bOTU22 *Veillonella dispar* and fOTU2 *Candida palmioleophila* were most significant (Table S3-4). In comparison, the same analysis yielded a reduced interaction network for healthy controls with 86 significant correlations, the majority of which (78, 90.7%) were B-B interactions (Fig. 4a-b). To adjust for sample size, we reconstructed disease network using a balanced sample size with healthy subjects (n=29). Despite a relatively smaller network, the network topology generally resembled that using all subjects (Fig. 4a, Fig. S7a), indicating sample size was likely not confounding the different networks between the two groups.

**Figure 4.**
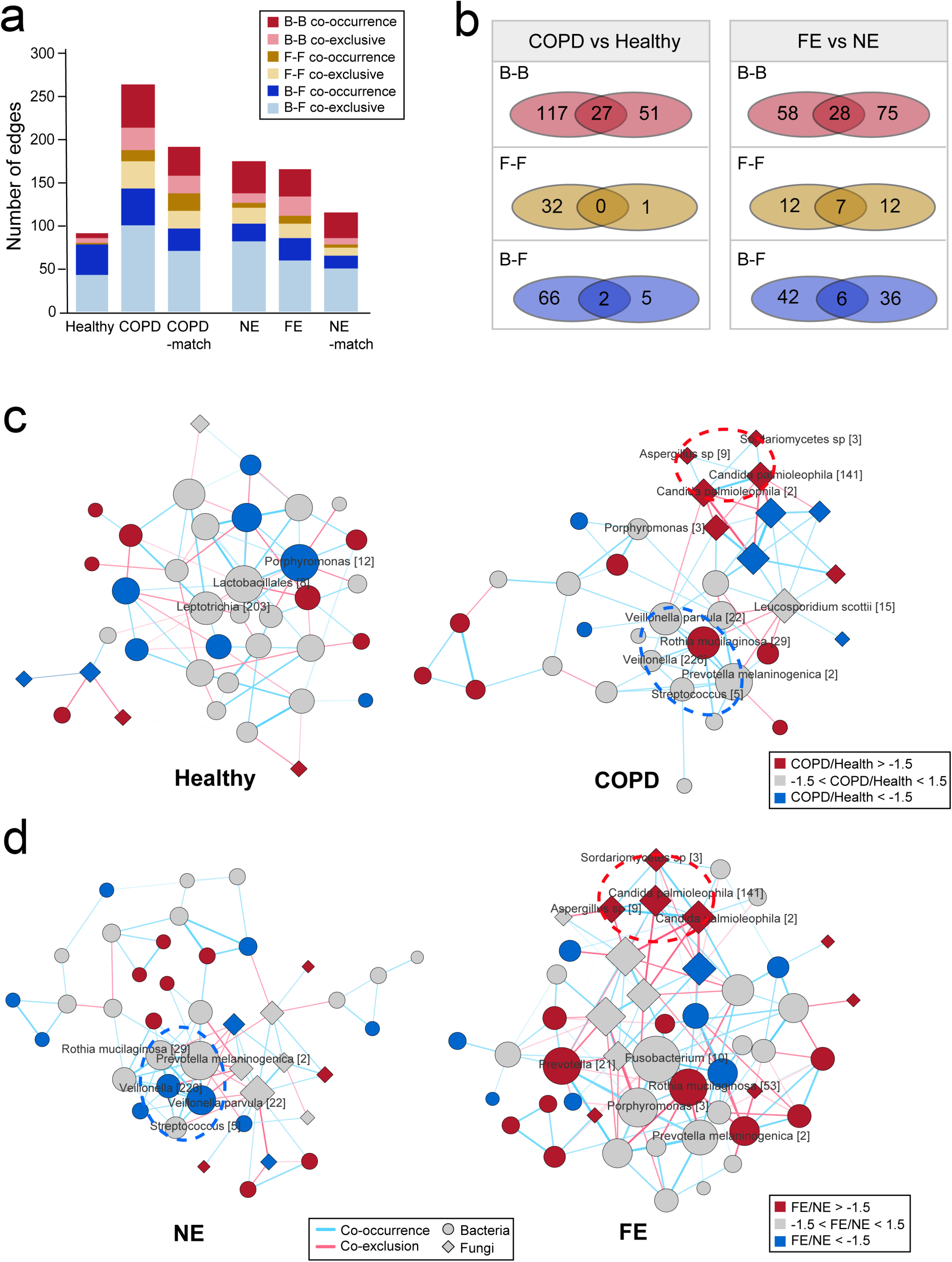
Airway bacterial and fungal interaction networks. **(a)** The number of significant bacterial-bacterial (B-B), bacterial-fungal (B-F) and fungal-fungal (F-F) interactions in the networks of healthy controls, COPD patients, the subgroup of COPD patients with healthy-matched sample size (COPD-match), non-frequent exacerbators (NE), frequent exacerbators (FE) and the subgroup of NE with FE-matched sample size (p<0.05). **(b)** Venn diagram for the shared and unique B-B, B-F and F-F interactions between COPD patients and healthy controls, and between FE and NE. **(c-d)** Bacterial and fungal interaction networks for healthy controls and COPD patients (c), and for NE and FE (d). Nodes were shaped by bacterial or fungal OTUs, and colored by their fold changes in COPD versus healthy groups or in FE versus NE groups. The size of the node is proportional to its degree of connectivity. Edges were colored by co-occurrence (blue) and co-exclusive (red) interactions. Edge width is proportional to the absolute correlation score. Only significant interactions with SparCC correlation>0.3 were shown for visualization purpose. Module 1 is highlighted in red dotted ellipse. Module 2 is highlighted in blue ellipse. The full interaction networks are in Fig. S6.

### Bacterial-fungal interactions in frequent and non-frequent exacerbators

We further performed sub-analysis on the interaction network for the FE and NE groups. Despite comparable network size, there were notable differences between bacterial and fungal interactions between the two groups (Fig. 4d, Fig. S6c-d). For example, in the FE group, four fOTUs, fOTU141 *Candida palmioleophila*, fOTU2 *Candida palmioleophila*, fOTU9 *Aspergillus* spp. and fOTU3 Sordariomycetes spp. showed strong mutual positive correlations in a subnetwork (module 1), which together exhibited co-exclusive relationships with most other fOTUs (Fig. 4c, Fig. S6b). This module was however absent in the network for NE group. On the other hand, another subnetwork, consisting of the mutual co-occurrence relationships between bOTU29 *Rothia mucilaginosa*, bOTU226 *Veillonella* spp., bOTU2 *Prevotella melaninogenica*, bOTU15 *Prevotella* spp. and bOTU5 *Streptococcus* spp. (module 2), was specifically present in the network for NE group. Again, network reconstruction using a balanced sample size (n=26) indicated that the different networks observed were not related to sample size (Fig. S7b). Network reconstruction using Spearman correlation mostly recapitulated the findings using SparCC (FDR p<0.2, Fig. S8), indicating the differences in interaction network were robust to the algorithm used.

### Correlations of bacterial and fungal microbiome with inflammatory mediators in COPD

To investigate interactions between bacterial and fungal microbiome and host response, we performed correlation analysis between bacterial, fungal OTUs and inflammatory mediators on a subset of 40 COPD patients with all data available. Fifty-nine correlations were found between blood or sputum mediators and bacterial or fungal OTUs (FDR p<0.20, Fig. 5). Of them, blood IL-6 was negatively correlated with bOTU8 Lactobacillales spp. and bOTU12 *Porphyromonas* spp. that were the hub OTUs in healthy controls, while it was positively correlated with fOTU8 *Cryptococcus* spp. that was present in the COPD network. Sputum IL-6 was negatively correlated with bOTU5 *Streptoccocus* spp. and bOTU29 *Rothia mucilaginosa* that was part of the module 1, and positively correlated with fOTU5 *Aspergillus* spp. Sputum IL-8 was positively correlated with fOTU3 Sordariomycetes spp., fOTU9 *Aspergillus* spp., fOTU15 *Leucosporidium scottii* and fOTU85 *Aspergillus penicillioides*, the former two being part of module 2 featured in the network for FE group. We were not able to perform the sub-analysis for frequent and non-frequent exacerbators separately due to the small number of patients in each subgroup with available mediator measurements.

**Figure 5.**
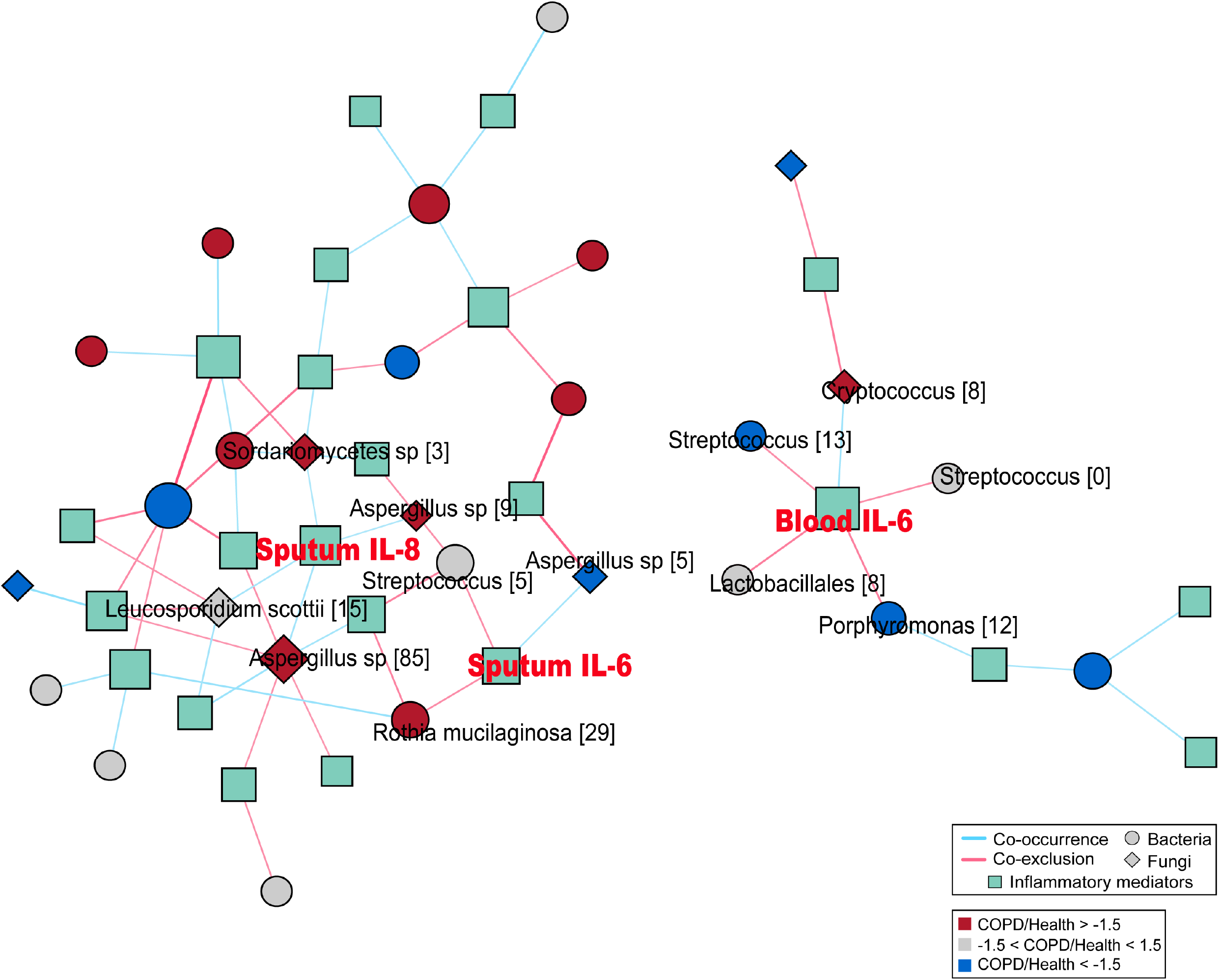
Interaction network between airway bacterial and fungal microbiome and host inflammatory mediators. Nodes were shaped by bacterial or fungal OTUs or inflammatory mediators. Bacterial and fungal OTUs were colored by their fold changes in COPD patients versus healthy controls. The size of the node is proportional to its degree of connectivity. Edges were colored by co-occurrence (blue) and co-exclusive (red) interactions. Edge width is proportional to the absolute correlation score in Spearman correlation (FDR p<0.20). Only correlations between bacterial/fungal OTUs and host inflammatory mediators were shown for visualization purpose.

## Discussion

Here we reported the first simultaneous characterization of the airway bacterial and fungal microbiome in COPD. The airway mycobiome was predominated by the phylum Ascomycota over Basidiomycota, consistent with observations in the airways of bronchiectasis and cystic fibrosis(17, 18, 38) but opposite to one study in asthma(15). We observed significant increases of pathogenic fungal taxa including *Candida*, *Cryptococcus* and *Schizophyllum* in COPD patients. Members of *Schizophyllum* and *Aspergillus* participate in invasive infections and provoke host immune recognition(39). *Cryptococcus* is known to interact with airway epithelium and lead to enhanced allergic inflammation(40). Overall there were greater community shifts for fungi than bacteria in COPD patients versus controls, and in FE versus NE. Supervised learning analysis identified a set of bacterial and fungal OTUs that together showed the optimal discriminatory potential for COPD patients and the frequent exacerbator phenotype, although cross-validation of these features in independent cohorts is warranted.

Importantly, there was a significant negative correlation between bacterial and fungal alpha diversity, both of which altered in opposite directions between COPD and healthy subjects and between FE and NE. Accordingly, individual bacterial and fungal OTUs showed disproportionately higher co-exclusive than co-occurrence relationships with each other. In particular, commensal bacterial taxa such as *Prevotella* and *Veillonella* exhibited inverse relationships with pathogenic fungal taxa such as *Candida palmioleophila* and *Aspergillus* spp. These results support the notion that there was a delicate balance between bacterial and fungal communities in the airways. The disruption of such community balance, characterized by the loss of commensal bacterial taxa and enriched pathogenic fungal taxa, is implicated in COPD pathogenesis(41, 42).

We observed distinct patterns of bacterial-fungal interactions both between COPD patients and healthy controls and between FE and NE. In particular, there was an enhanced and more sophisticated microbial interaction in COPD compared to healthy controls, which reflected a more active crosstalk between members of microbiome in response to altered local airway environments in disease. In healthy state, the airway microbiome was dominated by commensal bOTUs that mostly exhibited co-occurrence interactions. In COPD, additional B-F and F-F interactions were involved. While a higher number of F-F interactions were positive, a larger proportion of B-F interactions were negative, a finding that coincides with one recent study on the gut fungal microbiome in colorectal cancer(43). Thus the co-occurrence intrakingdom and co-exclusive interkingdom interactions may be a signature for disease-associated human microbiome in general. This is also in align with the opposite trend of changes between bacterial and fungal diversity, and suggests that disruption of normal bacterial communities may provide pathogenic fungi with a favorable condition for intra-fungi interaction in COPD. Furthermore, several pro-inflammatory mediators such as blood and sputum IL-6 and sputum IL-8 that are known to associate with lung microbiome(4), were negatively correlated with commensal bOTUs in the health-related network, and positively correlated with disease-associated pathogenic fOTUs. Thus the perturbation of ecological interactions in COPD was also associated with increased airway and systemic inflammations.

There were also important differences in bacterial-fungal interactions between FE and NE. The most remarkable difference was the disappearance of five mutually co-occurring commensal bOTUs (module 2) and emergence of four mutually co-occurring pathogenic fOTUs (module 1) in FE. Our results suggest that there was further airway dysbiosis in frequent exacerbators characterized by the displacement of commensal bacterial interactions by pathogenic fungal interactions, which was also associated with enhanced airway inflammation. The emergence of pathogenic fungi in particular *Candida palmioleophila* and *Aspergillus* spp. could be a marker for the frequent exacerbators that drives the greater microbial perturbation and inflammation, and together lead to the accelerated disease progression and increased vulnerability to subsequent exacerbations.

There are several caveats to our study. First, the study design is single-centred and cross-sectional. Further bacterial and fungal microbiome surveys preferably in cohorts with different demographic background is warranted to validate our findings. Second, targeted amplicon sequencing has insufficient resolution in species-level identification, in particular for the fungal population with a lack of well-characterized reference database(6). Despite the attempt to improve the fungal taxonomy assignment using phylotyping algorithm, the fungal taxa that can be assigned to genus or species level remain limited. Third, due to limited sputum available, inflammatory mediators were characterized only for a subset of patients and not for healthy subjects, which limits our ability to perform more detailed analysis on host-microbiome interactions between COPD and healthy subjects and within different patient subgroups.

## Interpretation

In summary, we characterized the collective airway bacterial and fungal microbiome in COPD. We showed that the disruption of airway community balance, characterized by the enriched pathogenic fungal taxa over commensal bacterial taxa, is implicated in COPD and associated with airway inflammation. The airway mycobiome is an important cofactor mediating pathogenic infection and airway inflammation, and should be taken into account when assessing the role of airway microbiome in COPD.

## Acknowledgements

Author contributions: HZ and ZW had full access to all of the data in the study and takes responsibility for the integrity of the data and the accuracy of the data analysis. HL, RC and ZW conceived and designed the study. NC, XT, ZHL, ZYL and JS coordinated the collection of sputum samples and clinical data. HL, FW, YY and CL processed the sputum samples, performed DNA extraction and library preparation. HL, YH and ZW performed all data analysis. RC and HZ supervised the study. ZW wrote the manuscript. All authors provided critical comments and approved the final version of the manuscript.

